# The intrinsically disordered AR2 domain of RNase E binds mRNA translation initiation regions

**DOI:** 10.64898/2026.08.11.744331

**Authors:** Daniel G. Mediati, Saleh Alquethamy, Christopher Jin, Jai J. Tree

## Abstract

Intrinsically disordered regions are widespread in RNA-processing machines. In *Escherichia coli*, RNase E uses its intrinsically disordered C-terminal domain (CTD) to recruit RNAs to the N-terminal catalytic domain, including mRNAs targeted by regulatory small RNAs (sRNAs), but the basis of substrate recognition and specificity is unclear. We engineered a protease-cleavable RNase E and used split-CRAC to isolate RNAs crosslinked to the AR2 sub-domain of the intrinsically disordered CTD fragment. AR2 preferentially engaged mRNAs and was depleted of sRNAs and sRNA-containing hybrids, supporting recognition of the mRNA. AR2 contacts concentrated on accessible A-rich motifs surrounding ribosome-binding sites and start codons, and purified AR2 recognised this motif *in vitro*. AR2 also contacted an AUAA motif in the *rne* translation-initiation region, and AR2 deletion increased RNase E abundance implicating this interaction in autoregulation. These findings define a relatively short AR2-binding motif and are consistent with CTD interactions with the 30S subunit that may provide additional specificity for a subset of mRNA translation initiation regions.

**SIGNIFICANCE STATEMENT:** Most RNA turnover in bacterial cells is carried out by the RNA degradosome, yet how this molecular machine checks and selects RNAs for degradation remains incompletely understood. We show that an intrinsically disordered region of the degradosome enzyme RNase E, termed AR2, preferentially binds A-rich sequences near sites of translation initiation. Through recognition of this shared sequence feature in a common functional context, AR2 may help the degradosome recognise messenger RNAs as a functional class. AR2 also contributes to feedback control of RNase E expression by recognising its own messenger RNA.

## INTRODUCTION

RNase E is the major endoribonuclease responsible for bulk RNA turnover and is the scaffold of the multi-protein RNA degradosome in *Escherichia coli* [1]. RNase E contains a conserved, structured N-terminal domain (NTD) and a less-conserved, intrinsically disordered C-terminal domain (CTD) that recruits canonical degradosome components that include PNPase, Enolase, and RhlB [2, 3]. The NTD provides several well-defined routes for substrate recognition: the catalytic centre binds and cleaves single-stranded AU-rich RNA [4], the 5′ sensor pocket recognises monophosphorylated RNA ends [5, 6], and the direct-entry site accommodates internal and structured substrates [7]. By contrast, how the intrinsically disordered CTD selects specific RNAs for delivery to the catalytic domain remains poorly defined.

The CTD contains two arginine-rich RNA-binding regions, RBD and AR2 [8–13], and is proposed to capture the RNA before its transfer to the catalytic core. RNA binding compacts the CTD towards the NTD, providing a potential route for substrate handover [14]. Multiple regulatory small RNA (sRNA) recruit RNase E to facilitate processing of target mRNAs, and the CTD contributes to degradation of these sRNA–mRNA duplexes [8, 12, 15–20]. Earlier models proposed that Hfq recruited the duplex to RNase E [21], with the sRNA mediating interactions with the CTD of RNase E [8]. More recent work has found that both the mRNA-binding surface of Hfq and AR2 are required for efficient recruitment, implicating the duplexed mRNA in recognition by RNase E [12]. However, the RNA features recognised by AR2, and how this recognition is coordinated with sRNA pairing remain unknown.

Recent work has found that RNase E can interact with 30S subunits [22], with mRNA in 30S pre-initiation complexes (PIC) [14, 19], and nascent transcripts within coupled transcription– translation complexes [23]. In these complexes, the mRNA translation-initiation region can be presented for sRNA pairing and cleavage by RNase E [19, 23]. Notably, an RNase E CTD segment containing AR2, but not RBD, is required for efficient cleavage of mRNAs associated with 30S subunits *in vitro* [19]. This finding suggests that the CTD of RNase E may act at an early checkpoint where an mRNA is presented to cognate sRNAs before it proceeds into translation.

Here, we looked to determine substrates of the AR2 domain within the intrinsically disordered CTD of RNase E. We engineered a protease-cleavable RNase E that releases an AR2-containing CTD fragment and used protease cleavable *in vivo* RNA crosslinking and analysis of cDNA (split-CRAC) to isolate RNAs crosslinked to the AR2 domain. AR2 preferentially engaged mRNAs and was depleted of sRNAs and sRNA-containing duplexes. We demonstrate that AR2 contacts are concentrated on accessible A-rich motifs within mRNA translation-initiation regions, and purified AR2 recognised this motif *in vitro*. These findings identify a relatively short recognition motif for the intrinsically disordered AR2 RNA-binding domain and suggests that the A-rich motif may achieve specificity through additional interactions with the translation initiation machinery. AR2 recognition also contributes to RNase E autoregulation by binding the *rne* translation-initiation region, demonstrating how this pathway can influence gene expression. Our data support a translation initiation surveillance model in which AR2 first engages an accessible mRNA initiation region before cognate sRNA pairing commits the transcript to decay.

## RESULTS

### The AR2-containing CTD fragment of RNase E preferentially binds mRNAs

The RNA sequences and structures recognised by the intrinsically disordered CTD of RNase E are poorly defined, in part because the two RNA-binding sites, RBD and AR2, lie within an unstructured region that has resisted the structural analysis applied to the catalytic NTD. To isolate RNAs engaged by the region containing AR2, we used split-CRAC, which allows protease cleavage and release of a defined portion of RNase E together with its UV-crosslinked RNA. Using CRISPR-Cas9, we introduced a TEV cleavage site at Val744 into the chromosomal *rne* CTD, 52 residues upstream of AR2, and appended a C-terminal His_6_-TEV-FLAG_3_ (HTF) tag (**Figure 1A**). TEV cleavage released a protein fragment containing AR2 together with the downstream Enolase- and PNPase-binding regions; we therefore refer to this species as the AR2-containing CTD fragment (AR2 for brevity). Dual-affinity purification followed by TEV cleavage and elution recovered this fragment, and we verified this by Western blot (**Figure 1Bi**).

**Figure 1.**
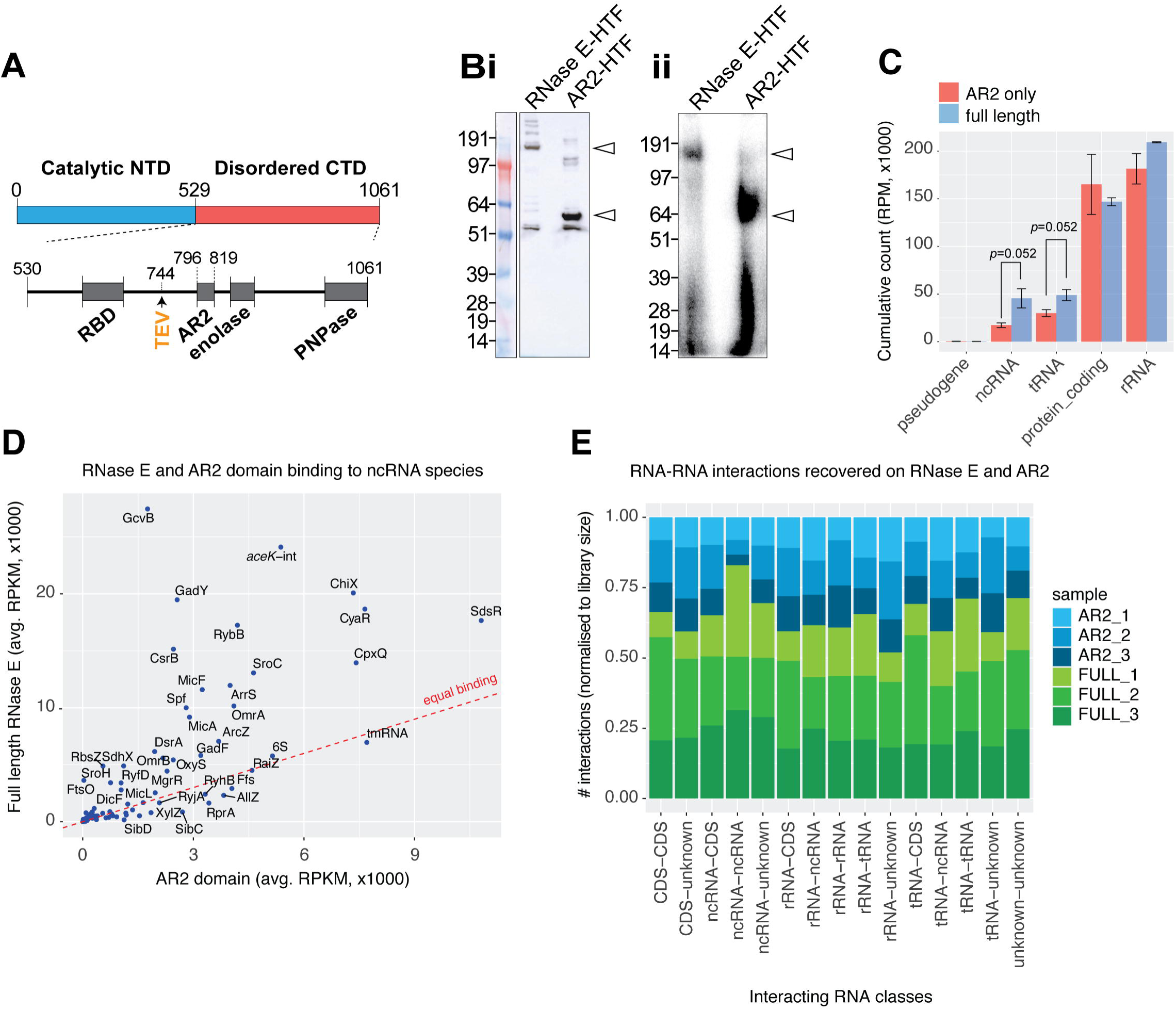
Split-CRAC isolates an AR2-containing CTD fragment depleted of small RNAs and RNA–RNA hybrids. (**A**) Schematic of *E. coli* RNase E showing the catalytic N-terminal domain (NTD), intrinsically disordered C-terminal domain (CTD), RNA-binding domain (RBD), AR2, and the enolase- and PNPase-binding regions. The engineered TEV cleavage site at Val744 releases the AR2-containing CTD fragment from RNase E-HTF. (**B**) Recovery of full-length RNase E-HTF and the AR2-containing fragment following dual-affinity purification and TEV elution. (**i**) SDS–PAGE analysis and (**ii**) autoradiogram of 5′-radiolabelled RNA–protein complexes. Molecular-mass markers (kDa) are shown at left; arrowheads indicate full-length RNase E and the AR2-containing fragment. (**C**) Cumulative reads recovered on full-length RNase E or the AR2-containing fragment for each RNA class. Reads were converted to reads per million (RPM) to normalise between libraries; error bars represent the standard error of three biological replicates. (**D**) Average abundance of individual non-coding RNAs in full-length RNase E and AR2-fragment libraries. The dashed line indicates equal recovery. (**E**) RNA–RNA interactions recovered in full-length RNase E and AR2-fragment libraries, grouped by the RNA classes in each hybrid and normalised to library size. Colours denote biological replicate libraries.

To map the RNAs bound by AR2 across the transcriptome, UV-crosslinked complexes from the AR2-HTF strain, a full-length RNase E-HTF strain, and an untagged control were purified under denaturing conditions, 5′-radiolabelled, and processed for CRAC sequencing (**Figure 1Bii**) [24]. Coding sequences were recovered on AR2 at similar or slightly higher levels than on full-length RNase E, whereas non-coding RNAs—including sRNAs—and tRNAs showed a trend towards depletion on AR2 (*p* = 0.052; **Figure 1C**). Depletion of tRNAs is consistent with their recognition by the NTD direct-entry site rather than the CTD [25]. Plotting individual non-coding RNAs confirmed that almost all sRNAs were depleted on AR2 relative to full-length RNase E (**Figure 1D**).

Because the CTD is thought to recruit sRNA–mRNA duplexes, we asked whether AR2 engages the sRNA within these duplexes by examining RNA–RNA hybrids recovered from each sample. Despite the larger AR2 libraries, hybrid recovery was reduced on AR2, and interactions involving non-coding RNAs—both ncRNA–ncRNA and ncRNA–mRNA—were strongly depleted (Figure 1E). Together, these data show that the AR2-containing fragment is depleted of small RNAs and their duplexes. This pattern argues against direct or stable recognition of the sRNA component by AR2 and instead supports preferential engagement of the mRNA component.

### AR2 is enriched at mRNA translation-initiation regions

To determine where AR2 binds on transcripts, we constructed metagene plots of cumulative read density centred on the start or end of features from each class of RNA. Consistent with the class-level depletion described above, AR2 recovery was low for sRNAs and tRNAs (**SI Figure 1**). Within mRNAs, AR2 was depleted at both the 5’ and 3′UTRs but recovered at similar or slightly higher levels than full-length RNase E at the start and stop codons (**Figure 2A**), demonstrating that AR2 binding concentrates at the boundaries of open reading frames rather than at UTR ends.

**Figure 2.**
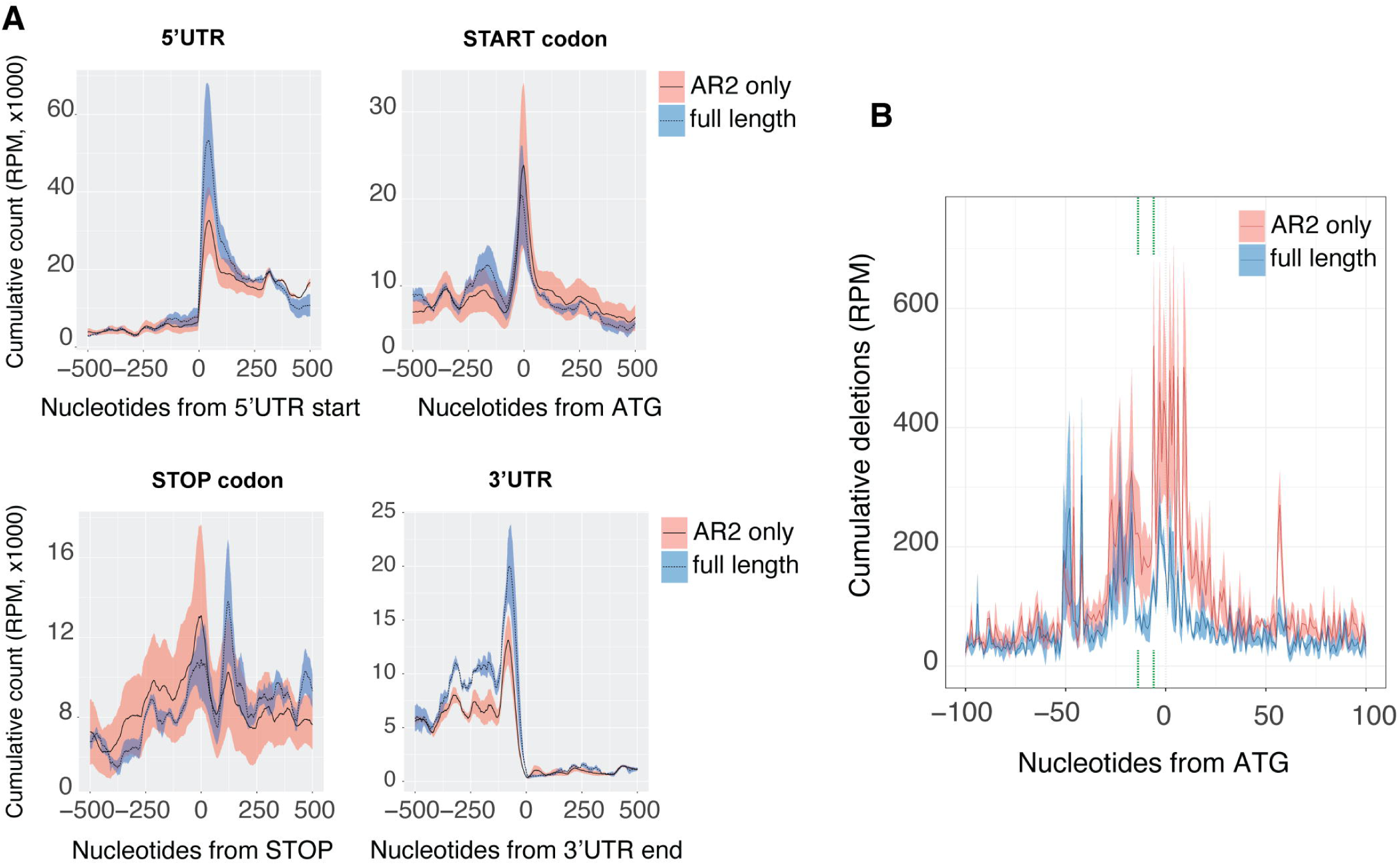
AR2 is enriched at mRNA translation-initiation regions. (**A**) Cumulative read density for full-length RNase E (blue) and the AR2-containing fragment (red), mapped relative to 5′UTR starts, start codons, stop codons and 3′UTR ends. Lines represent the mean of three biological replicates and shaded intervals represent standard deviation. (**B**) Contact-dependent deletions mapped relative to mRNA start codons for full-length RNase E (blue) and the AR2-containing fragment (red). Lines represent the mean of three biological replicates and shaded intervals represent standard deviation. Dashed grey lines indicate the approximate position of the ribosomal binding site.

To localise these contacts at nucleotide resolution, we mapped deletions introduced at UV-crosslinking sites, which mark positions of protein contact. Within mRNAs, deletions on AR2 were maximal across a window extending from immediately 5′ of the ribosome-binding site (RBS) to the 3′ side of the start codon but were reduced directly over the RBS itself (**Figure 2B**). This pattern indicates that AR2 contacts sequences flanking the RBS and start codon while either avoiding, or being occluded from, the RBS. Although metagene coverage was elevated at both open-reading-frame boundaries, the strongest nucleotide-resolution signature of AR2 binding occurred at the translation-initiation region.

### AR2 recognises an A-rich motif within translation-initiation regions

To identify the sequence recognised by AR2, we extracted high-confidence binding sites defined by contact-dependent deletions (in >10 reads) and examined nucleotide frequency at the point of crosslinking. AR2 contact sites were strongly enriched for adenosine and defined an AUAA motif (**Figure 3A**), with maximal crosslinking at the first adenosine. Motif analysis of the flanking sequence with MEME-ChIP [26] recovered a significant A-rich motif that contained the AUAA sequence and could accommodate a start codon (E-value = 2.7 × 10^−10^; **Figure 3B**). Mapping AR2 reads onto all genomic matches identified with FIMO [27] confirmed enrichment of AR2 binding at the motif across the genome, rather than only within the subset of highly crosslinked mRNAs (**Figure 3C**).

**Figure 3.**
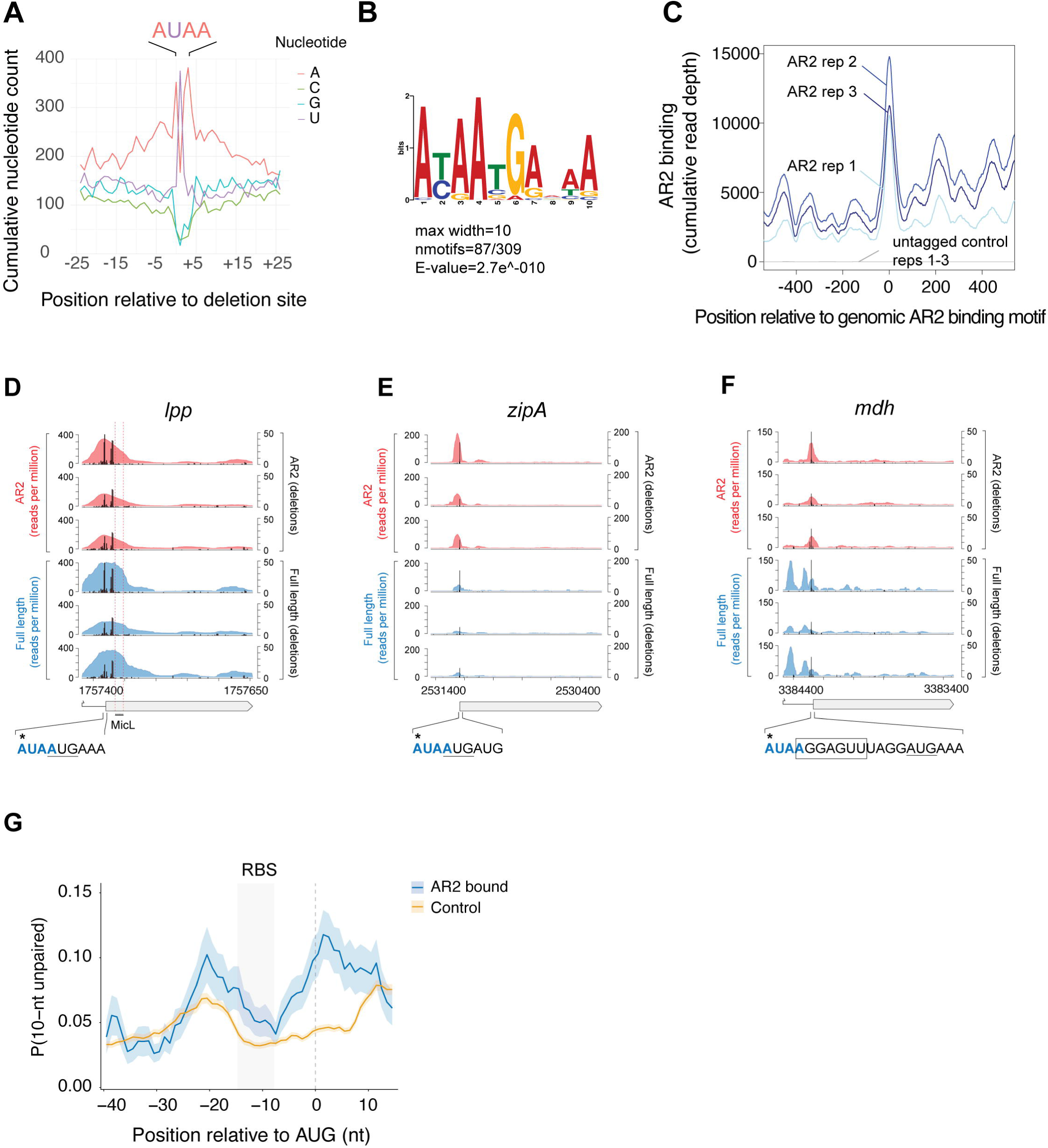
AR2 recognises an accessible A-rich motif within mRNA translation-initiation regions. (**A**) Nucleotide frequency surrounding contact-dependent deletion sites within high-confidence AR2 binding sites. (**B**) A-rich motif recovered from AR2 binding sites by MEME-ChIP. (**C**) AR2 read density around genomic matches to the motif identified by FIMO (*p* < 0.0001). Traces show three AR2 biological replicates and three untagged-control replicates. (D–F) Binding of the AR2-containing fragment (red) and full-length RNase E (blue) to *lpp* (**D**), *zipA* (**E**) and *mdh* (**F**). Filled profiles show read coverage and black vertical lines show contact-dependent deletions. The CDS and 5′ UTR are shown below each track. Expanded sequences indicate the AR2 motif (blue), start codon (underlined), predicted RBS (boxed where shown) and position of maximal deletion (*). (**G**) Predicted RNA accessibility, expressed as the probability of a 10-nt window being unpaired, around the start codon of AR2-bound mRNAs and matched control mRNAs. The predicted RBS interval is shaded.

We next assessed enrichment of the AR2 motif at mRNA translation-initiation regions. Inspection of individual transcripts supported AR2 contact at AUAA motifs overlapping or flanking translation-initiation regions. In *lpp*, encoding the outer-membrane lipoprotein Lpp, and *zipA*, encoding an essential cell-division protein, maximal AR2 deletions fell within an AUAA motif overlapping the start codon (**Figure 3D&E**). In *mdh*, encoding malate dehydrogenase, full-length RNase E made distributed contacts across the 5′UTR, whereas AR2 binding concentrated on an AUAA motif overlapping the predicted RBS upstream of the start codon (**Figure 3F**).

To test directly whether AR2 recognises this motif, and to attribute recognition to AR2 rather than to another component of the larger CTD fragment isolated by split-CRAC, we performed gel-shift assays with purified protein and a 66-nt *in vitro* transcribed *lpp* fragment spanning the 5′UTR and translation-initiation region. Following earlier work that quantified AR2 binding to tRNA^Phe^ [8], we purified the AR2 domain alone and in complex with RhlB and Enolase. The AR2–RhlB–Enolase complex bound the *lpp* fragment with high affinity (K_d_ = 72 nM), and isolated AR2 also bound, albeit weaker (K_d_ = 463 nM; **Figure 4A-E**). Mutating the AUAA motif of *lpp* to GGGGG reduced AR2 binding (K_d_ = 865 nM; **Figure 4C&F**), supporting direct recognition of the A-rich sequence identified *in vivo*.

**Figure 4.**
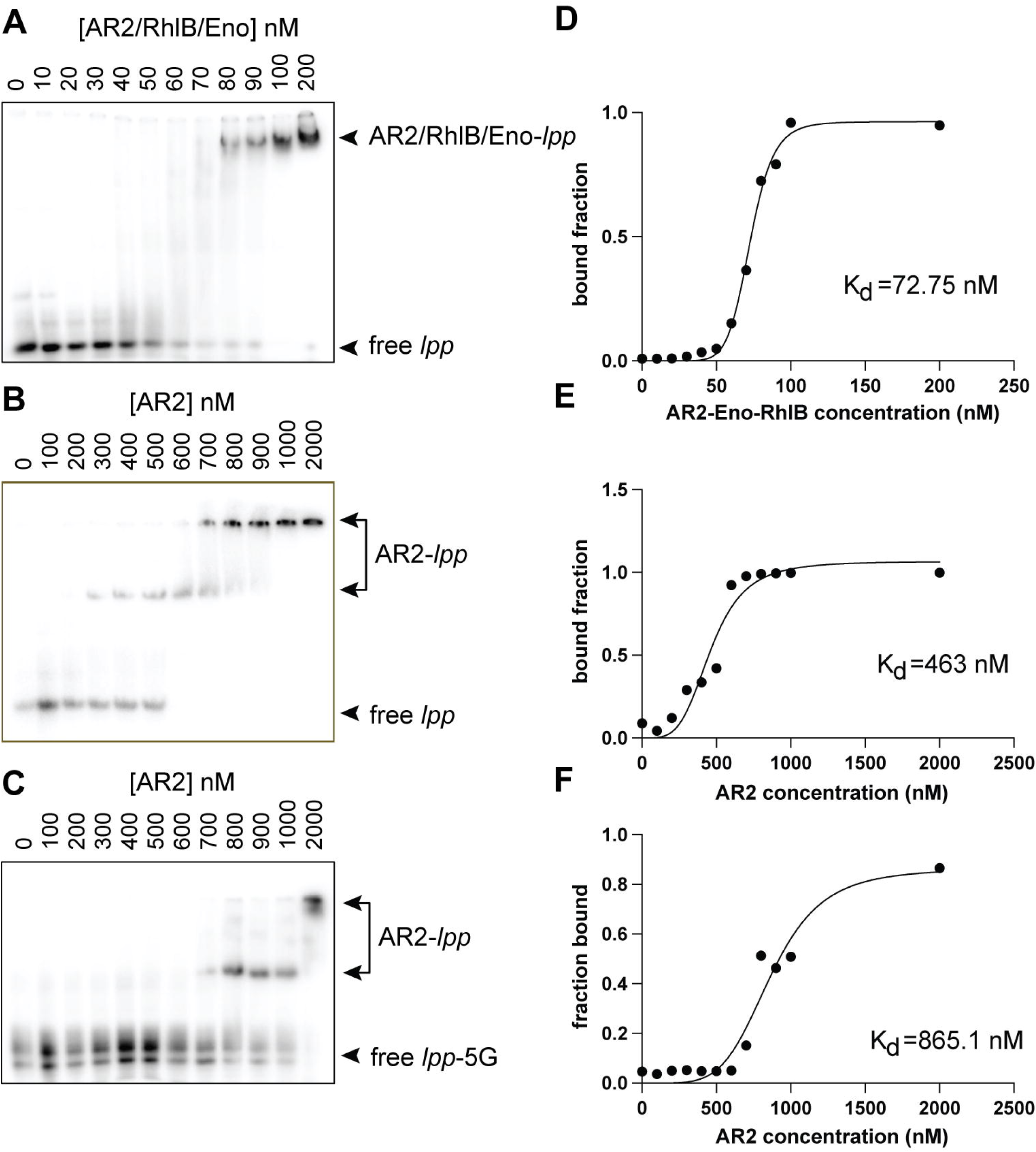
AR2 directly binds the lpp translation-initiation region. (**A**–**C**) Electrophoretic mobility-shift assays using the *lpp* translation-initiation-region RNA and increasing concentrations of the AR2–RhlB–enolase complex (**A**), isolated AR2 (**B**), or isolated AR2 with the AUAA-mutant *lpp* RNA (*lpp*-5G; **C**). Protein concentrations are indicated above the lanes; free and protein-bound RNA species are marked. (**D**–**F**) Bound fractions and fitted binding curves corresponding to panels A–C. Estimated dissociation constants (K_d_) are shown.

### AR2 crosslinking recovers abundant mRNAs with accessible translation-initiation regions

To characterise the mRNAs selected by AR2, we defined a high confidence target gene set as transcripts that carry AR2 contact-dependent deletions (>10 deletions/nt) within 20 nt of the start codon in at least 1 replicate, and a statistically significant match to the A-rich binding motif (FDR<0.001) (107 genes retained, Methods). This target gene set was not enriched for any functional category, for membrane-localisation, or for signal-recognition-particle-associated proteins (**SI Figure 2**), suggesting that AR2 selectivity reflects local features of the initiation region rather than a particular group of genes that share ontology or localisation. We next asked whether AR2-interacting mRNAs differed in stability or expression by interrogating publicly available datasets. The AR2 crosslinked gene set did not have significantly different mRNA stability, but were associated with higher mRNA and protein abundance, and higher protein synthesis rates (**SI Figure 4A-D**) [28–30]. The increase in mRNA and protein abundances was not seen for mRNAs that only encode the A-rich motif at start codons, indicating that additional features contribute to AR2 selection.

We next examined features encoded within the translation initiation region. The AR2 crosslinked mRNAs had slightly longer 5’ UTRs but did not have significantly different predicted translation initiation rates (**SI Figure 3A&B**). The AR2 crosslinked gene set were enriched for adenosine around the RBS and start codon (FDR=4.6×10^-9^; **SI Figure 3C)**, consistent with the presence of an A-rich motif and reminiscent of S1 binding sites that enhance translation at weak or structured translation initiation sites. To explore this further, RNA accessibility (defined as the probability of being unpaired) was predicted for the AR2 crosslinked mRNAs. AR2 target mRNAs were more accessible than a matched control set, particularly across the RBS (*p* = 0.058) and start codon (*p* = 5.11×10^-5^; **Figure 3G**), indicating that AR2 often engages its A-rich motif when the initiation region is presented in a single-stranded, accessible form.

### AR2 contributes to autoregulation of RNase E

Expression of *rne* is negatively autoregulated through RNase E recognition of its own 5′UTR, coupling RNase E abundance to the stability of its transcript [31, 32]. AR2 was enriched at the *rne* 5′UTR, where its contacts concentrated on an AUAA motif 5′ of the RBS. By contrast, full-length RNase E contacted the 5′UTR more broadly, consistent with NTD engagement of the regulatory stem-loops hp2 and hp3 [31, 32] (**Figure 5A**). This indicates that AR2 contacts the *rne* translation-initiation region through the same accessible A-rich motif that it recognises across the transcriptome.

**Figure 5.**
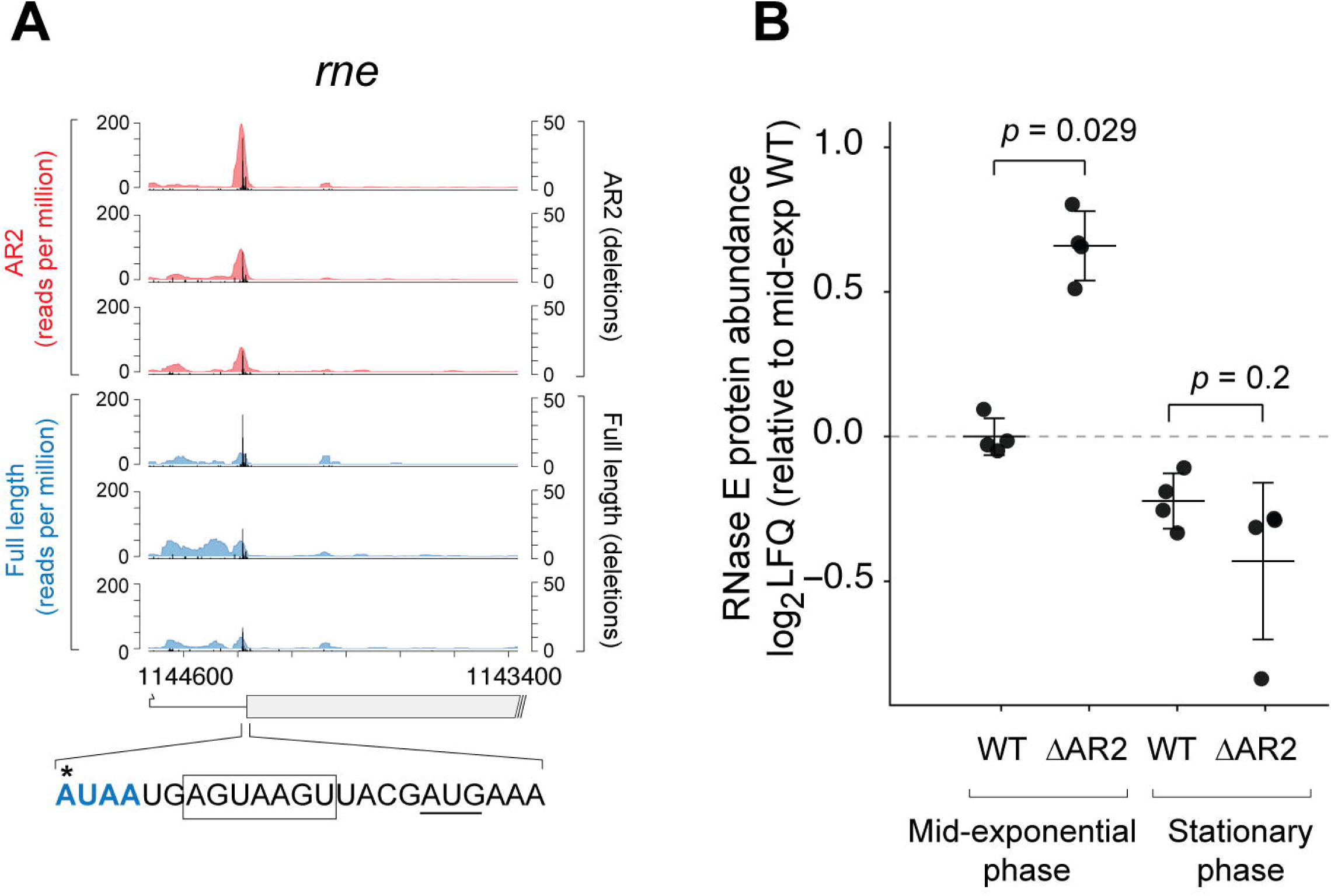
AR2 contributes to autoregulation of RNase E. (**A**) Binding of the AR2-containing fragment (red) and full-length RNase E (blue) across the *rne* translation-initiation region. Filled profiles show read coverage and black vertical lines show contact-dependent deletions. The expanded sequence indicates the AR2 motif (blue), predicted RBS (boxed) and position of maximal deletion (*). (**B**) RNase E protein abundance in wild-type (WT) and ΔAR2 cells during mid-exponential and stationary phase, shown as log2 fold change relative to mid-exponential WT. Points represent four biological replicates; horizontal lines and error bars show the mean and standard deviation. *p*-values were calculated using a Mann-Whitney U test.

To ask whether AR2 contributes to this autoregulation, we deleted the AR2 domain from the chromosomal copy of *rne* (ΔAR2) using CRISPR-Cas9 and compared the proteomes of ΔAR2 and its isogenic parent by label-free LC-MS/MS during mid-exponential and stationary growth phase. Relatively few AR2 target proteins changed significantly, and those that did showed no coordinated direction of change, indicating that loss of AR2 did not co-ordinately disrupt expression of its direct targets (**SI Figure 5**). However, RNase E itself was among the differentially abundant AR2 targets, potentially explaining the pleiotropic changes in the proteome. Its abundance increased by 47.4% (*p*=0.029) in the ΔAR2 strain during mid-exponential growth phase, consistent with impaired autoregulation when RNA turnover is most active (**Figure 5B**).

The direct contact of AR2 with the *rne* initiation region, together with increased RNase E abundance following deletion of the AR2 domain, supports a functional contribution of AR2 to RNase E autoregulation.

## DISCUSSION

RNase E recognises RNA through several sites with distinct specificities. Within the catalytic NTD, the catalytic centre cleaves single-stranded A-rich sequences [4], the 5′ sensor pocket binds monophosphorylated 5′ ends [7], and the direct-entry site accommodates structured RNA stems [25]. Our data identified a fourth specificity contributed by the intrinsically disordered C-terminal AR2 domain. Using split-CRAC, we find that AR2 preferentially engages an accessible A-rich motif within the translation-initiation region of mRNAs while being relatively depleted of sRNAs and tRNAs. These findings define an additional pathway through which the disordered CTD contributes to substrate selection by RNase E.

The relatively short AUAA recognition motif, and weaker interactions between purified AR2 and *lpp* mRNA *in vitro*, suggests that AR2 may use additional features to achieve specificity for the translation-initiation region. AR2 crosslinked mRNAs are more accessible at the RBS and start codon which may partly contribute to specificity. Depletion of AR2 contact-dependent deletions around the RBS and start codon also suggests that AR2 may be occluded from the RBS during interactions with mRNA translation-initiation regions. Recent *in vitro* work has found that mRNA initiation regions can be presented to RNase E for cleavage within the 30S pre-initiation complex (PIC) and the early expressome [19, 23]. The CTD recognition core, which includes AR2, is required for efficient mRNA cleavage in the PIC [19, 23] and can also form stable complexes with the 30S subunit [22]. These data suggest that AR2 depletion at the RBS could be due to occlusion by a 30S subunit. We propose that CTD interactions with the 30S subunit may contribute to AR2 specificity for the translation initiation region. This composite recognition model would explain how a short motif acquires specificity for translation-initiation regions, although direct testing will be required.

This model may also help explain why AR2-interacting mRNAs have increased translation and protein abundance. RNase E may sample accessible initiation regions without necessarily committing every bound transcript to decay. This is consistent with a model of RNase E CTD interactions at an early checkpoint in translation initiation [19, 23], with degradation requiring or accelerated by an additional signal such as sRNA pairing or engagement of the NTD.

The CTD contributes to the degradation of several sRNA–mRNA duplexes [8, 12, 15–20], and sRNA–mRNA pairing has been proposed to recruit these duplexes to RNase E through AR2 [8, 12]. We instead find that AR2 is depleted of sRNAs and sRNA-mRNA duplexes, and preferentially engages mRNAs. These observations support a model in which AR2 recognises the mRNA component of a regulatory complex [12]. Because AR2 contacts often overlap the translation-initiation regions used by many sRNAs, the data suggest that AR2 engages mRNAs before sRNA pairing and NTD-dependent cleavage. Interestingly, the unpaired AUAA motif fits broadly with the recognition motif of the NTD catalytic site that is enriched at start and stop codons [4].

A functional consequence of AR2 recognition is suggested by *rne* itself. RNase E autoregulates its synthesis through *cis*-acting elements in the *rne* 5′UTR, including the structured stem-loops hp2 and hp3 recognised by the catalytic NTD [31, 32]. We find that AR2 contacts an AUAA motif 5′ of the *rne* RBS that is unstructured and deletion of AR2 significantly increases RNase E abundance during mid-exponential growth. AR2 may therefore provide a CTD-mediated contribution to a feedback loop previously attributed principally to the NTD catalytic domain, potentially by recruiting the accessible initiation region for NTD-dependent recognition.

Intrinsically disordered regions are widespread in RNA-processing proteins, yet how they contribute to selective RNA recognition remains incompletely understood. Our findings show that the disordered AR2 domain of RNase E preferentially engages an accessible A-rich motif within mRNA translation-initiation regions. The short motif is unlikely to confer this specificity alone; instead, recognition may depend on the combined presentation of the RNA motif within the 30S pre-initiation complex. The combination of CTD interactions with the 30S subunit and AR2 interactions with a short motif presented in this context may confer specificity for a subset of mRNAs at a translation initiation checkpoint.

## METHODS

### Bacterial strains and strain construction

Bacterial strains, plasmids and oligonucleotides are listed in Supplementary Tables 1–3. *Escherichia coli* MG1655 derivatives carrying a C-terminal His –TEV–3×FLAG tag on RNase E, an additional TEV site at Val744, or deletion of the AR2-containing CTD segment were constructed using the pCas/pTarget CRISPR–Cas9 system [33]. All modifications were verified by PCR and loss of the editing plasmids was confirmed. Culture and selection conditions are detailed in Supplementary Methods.

### Split-CRAC and CLASH

Split-CRAC was adapted from established E. coli CLASH and RNase III–CLASH workflows [34–36]. Untagged MG1655, RNase E–HTF and AR2–HTF cultures were analysed in three biological replicates. Cells were UV-C crosslinked, RNase E complexes were isolated by sequential FLAG and Ni–NTA affinity purification with intervening TEV cleavage, associated RNAs were partially digested, radiolabelled and ligated to sequencing adaptors, and cDNA libraries were sequenced as 150-bp paired-end reads. Full purification and library-construction conditions are provided in Supplementary Methods.

### Sequencing data and motif analysis

CRAC and hybrid reads were processed with Hyb-CRAC-R [35], incorporating pyCRAC [37] and *hyb* [38], and mapped to the *E. coli* MG1655 genome (NC_000913.3). Transcript features were obtained from RegulonDB [39]. Contact-dependent deletions were used to define direct AR2 contacts. Motifs were identified using MEME-ChIP [26], genomic matches were detected using FIMO [27], and overlapping sites were consolidated with BEDTools [40]. Normalisation, metagene, target-set and motif-analysis details are provided in Supplementary Methods.

### EMSA assays

The AR2 domain and AR2–RhlB–enolase complex were purified as previously described [8]. Binding of isolated AR2 or the AR2–RhlB–enolase complex to wild-type and AUAA-mutant *lpp* translation-initiation-region RNAs was quantified by electrophoretic mobility-shift assays. Detailed purification, immunoblotting and binding-analysis procedures are provided in Supplementary Methods.

### Quantitative proteomics

Wild-type and ΔAR2 cells were compared by label-free LC– MS/MS during mid-exponential and stationary growth using four biological replicates per genotype and phase. RNase E abundance was calculated from sample-wise median-normalised log LFQ intensities and compared within each growth phase using two-sided Mann–Whitney U tests. Sample processing, acquisition parameters and complete computational filtering are described in Supplementary Methods.

### Statistical analysis and data availability

Statistical tests, effect sizes and multiple-testing procedures are specified in the relevant figure legends and Supplementary Methods. Split-CRAC sequencing data are available from NCBI GEO under accession GSE317719 (reviewer token: cnonqmeihxujdcx). Mass spectrometry proteomics data generated in this study are available at the ProteomeXchange Consortium via the PRIDE partner repository with the dataset identifier PXD082328 (reviewer token: bU6SLAwOClFt).

## Acknowledgements

DGM was supported by the award of a Chancellor’s Research Fellowship from the University of Technology Sydney (UTS). JJT and DGM were supported by ARC Discovery grant (DP220101938). JJT, CJ and SA were supported by NHMRC Ideas grant (GNT2028572).

