## Supplementary Figures for "The intrinsically disordered AR2 domain of RNase E binds mRNA translation initiation regions"

**Supplementary Figure 1**

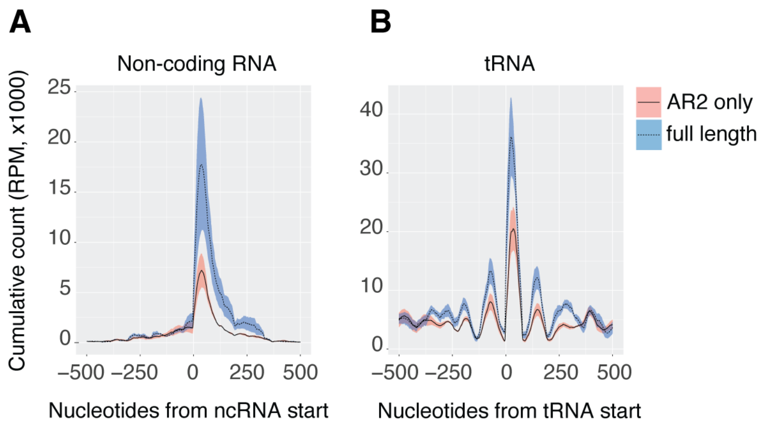


**SI Figure 1. AR2 interactions with ncRNA and tRNA**.

Metagene plots of the full-length RNase E (blue) and the AR2 containing CTD fragment of RNase E (red) binding to non-coding RNA (**A**) and tRNA (**B**). Lines represent the mean of three biological replicates and shaded intervals represent standard deviation.

**Supplementary Figure 2**

**A**

| GO Term | Study # | Study Freq. | Pop. Freq. | *p*-value | *q*-value | name | gene products |
|---|---|---|---|---|---|---|---|
| GO:0045900 | 2 | 4.1% | 0.06% | 2.2E-4 | 0.0679 | negative regulation of translational elongation | raiA,ettA |

**B**

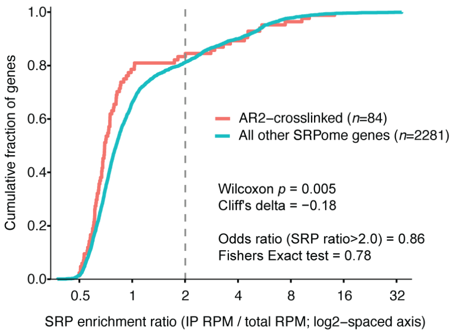


**C**

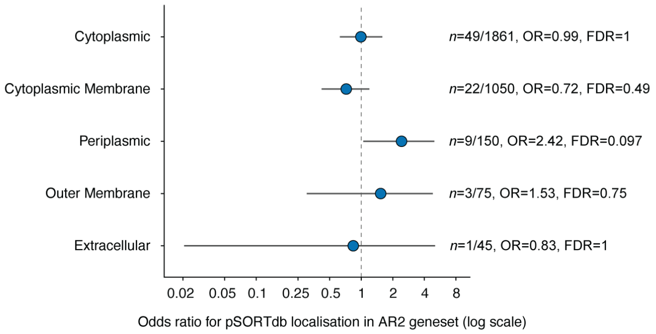


**SI Figure 2. Functional characteristics and predicted subcellular localisation of AR2-crosslinked genes.** (**A**) Gene Ontology enrichment analysis was performed using GOEnrichment within the Galaxy platform (1, 2). The highest-ranking Biological Process term was "negative regulation of translational elongation" (GO:0045900). No enriched terms were identified in the Cellular Component or Molecular Function ontologies. (**B**) Empirical cumulative distribution functions comparing SRP-interaction ratios for AR2-crosslinked genes ($n$ = 84) and all other genes represented in the published *E. coli* SRP interactome ("SRPome"; $n$ = 2,281) (3). The dashed vertical line indicates the SRP-interaction threshold of 2.0. The AR2 crosslinked geneset was modestly reduced for SRP interactions (Wilcoxon rank-sum test, $P$ = 0.005; Cliff's $\delta = -0.18$), and classification using the ratio >2.0 threshold provided no evidence of enrichment among AR2-crosslinked genes (odds ratio = 0.86; Fisher's exact test, $P$ = 0.78). (**C**) Enrichment of predicted protein-localisation categories among AR2-crosslinked genes was evaluated using PSORTb 3.0 predictions obtained from cPSORTdb for *E. coli* K-12 MG1655 (4, 5). Points show odds ratios and horizontal lines indicate 95% confidence intervals; the dashed line denotes an odds ratio of 1. $P$ values from category-specific Fisher's exact tests were adjusted for multiple testing using the Benjamini–Hochberg procedure. Periplasmic proteins showed the strongest nominal enrichment (odds ratio = 2.42; $P$ = 0.020), but this did not remain significant after correction ($q$ = 0.098).

### Supplementary Figure 3

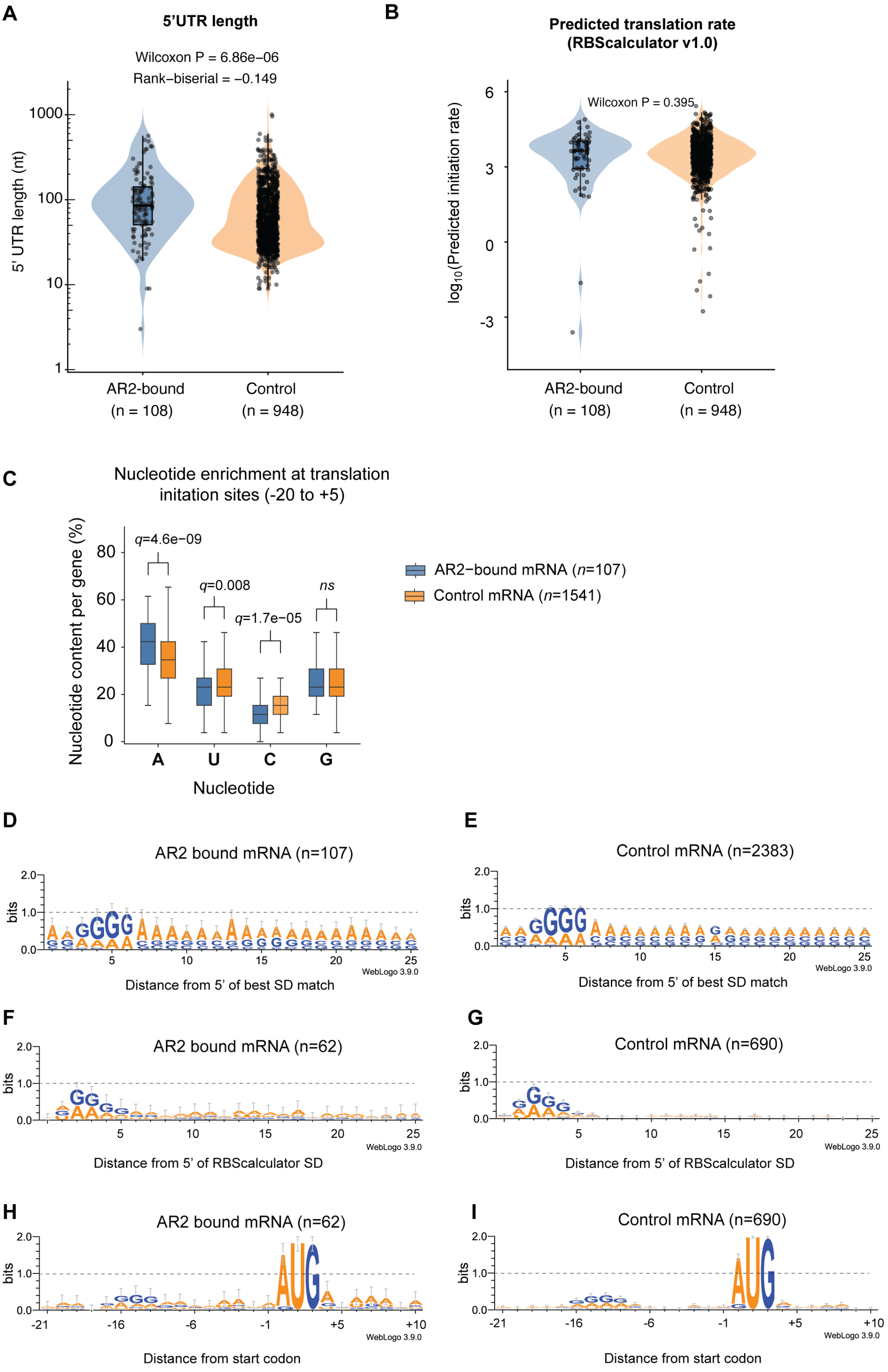

**SI Figure 3. Translation-initiation features of AR2-crosslinked mRNAs.**

(**A**) Distribution of annotated 5′-UTR lengths for AR2-crosslinked and first-in-operon control transcripts. All distinct 5′-UTR annotations in RegulonDB were retained, including alternative leaders; sample sizes therefore represent annotated UTR records (AR2, $n = 108$; control, $n = 948$), rather than unique genes (6). The y-axis is logarithmic. Violin plots show the distributions, internal boxes indicate the median and interquartile range, and points represent individual observations. Groups were compared using a two-sided Wilcoxon rank-sum test, with rank-biserial correlation reported as an effect-size measure. (**B**) Predicted translation-initiation rates calculated with RBS Calculator v1.0 (7) for unique genes with successful predictions (AR2, $n = 62$; first-in-operon control, $n = 690$). Values are displayed on a $\log_{10}$ scale and were compared using a two-sided Wilcoxon rank-sum test. Violin plots, boxes and points are defined as in A. (**C**) Per-gene nucleotide content in the 26-nt region from −20 to +5 relative to the first nucleotide of the annotated start codon for AR2-crosslinked mRNAs ($n = 107$) and the non-overlapping first-in-operon control set ($n = 1,541$). Boxes indicate the median and interquartile range, and whiskers extend to 1.5 times the interquartile range. AR2 and control distributions were compared separately for A, U, C and G using two-sided Wilcoxon rank-sum tests, with Benjamini–Hochberg correction across the four comparisons. AR2-crosslinked mRNAs had higher A content ($q = 4.6 \times 10^{-9}$) and lower U ($q = 0.008$) and C ($q = 1.7 \times 10^{-5}$) content; G content was not significantly different. (**D** and **E**) Sequence logos for AR2-crosslinked mRNAs (D; $n = 107$) and all non-AR2 control mRNAs (E; $n = 2,383$), aligned to the 5′ boundary of the best predicted Shine–Dalgarno match. (**F** and **G**) Sequence logos for the subset of AR2-crosslinked mRNAs (F; $n = 62$) and first-in-operon control mRNAs (G; $n = 690$) with a Shine–Dalgarno sequence predicted by RBS Calculator, aligned to the 5′ boundary of the predicted mRNA–16S rRNA pairing span. (**H** and **I**) The same AR2 and control subsets as F and G, aligned to the annotated translation start codon. Sequence logos in D–I were generated using WebLogo 3.9.0 (8). Total stack height indicates positional information content in bits, and relative letter height indicates nucleotide frequency at each position.

#### Supplementary Figure 4

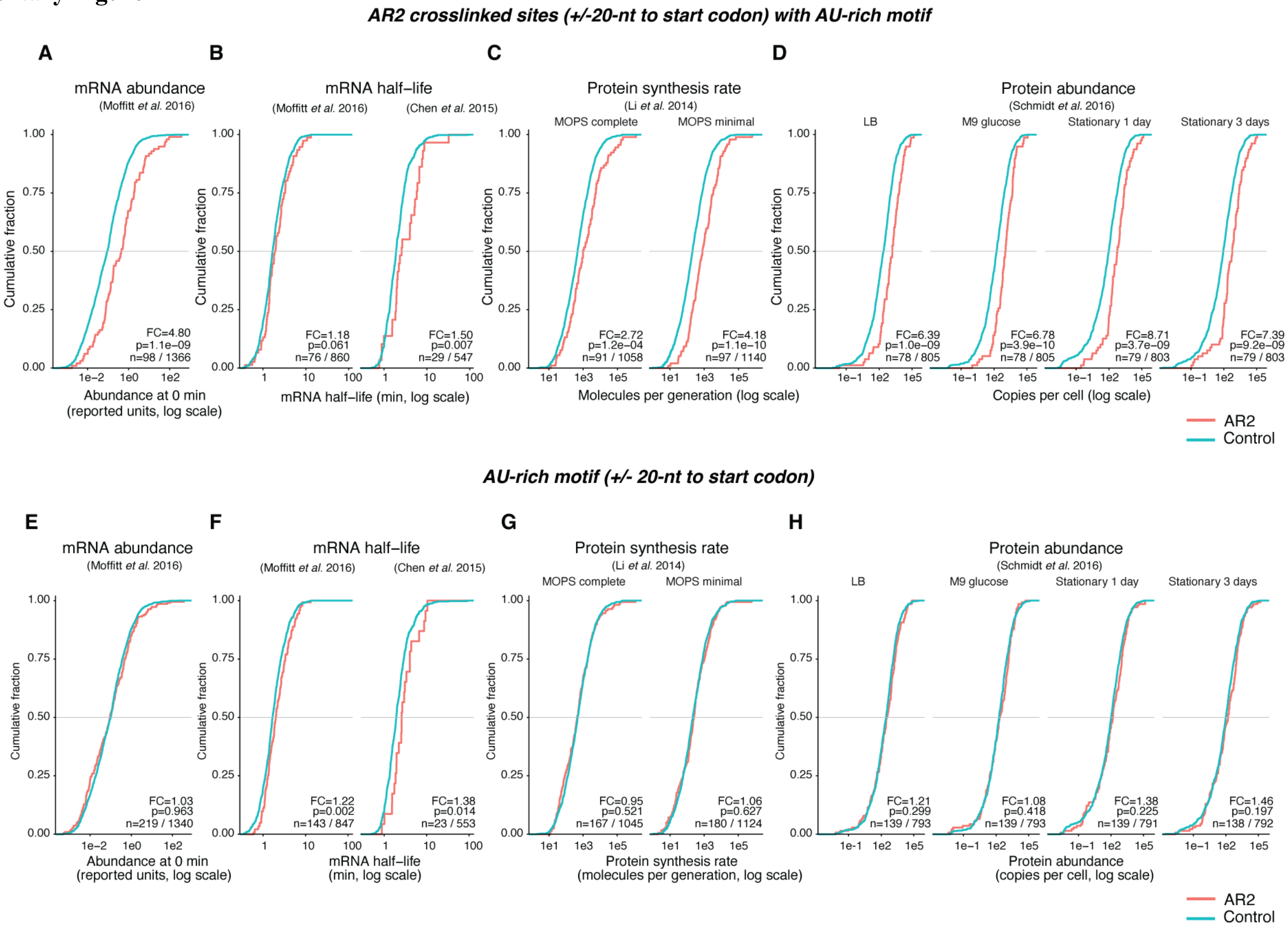

**SI Figure 4. Expression and stability characteristics of AR2-crosslinked and AR2-motif-containing mRNAs.**

Empirical cumulative distribution functions compare gene-expression measurements for AR2-associated gene sets (coral) with non-overlapping first-in-operon control genes (cyan). (**A–D**) Comparisons using the experimentally identified set of 107 AR2-crosslinked genes. (**E–H**) Comparisons using an alternative sequence-defined set of 250 genes containing a significant AU-rich AR2 motif within ±20 nt of the annotated start codon. For each row, genes belonging to the corresponding AR2 set were removed from the first-in-operon control set before matching to the available measurements; missing measurements were not imputed. (**A** and **E**) mRNA abundance at 0 min, averaged across untreated wild-type replicates, from Moffitt et al. (9). (**B** and **F**) mRNA half-lives determined independently by Moffitt et al. (9) and Chen et al. (10). For the Moffitt dataset, half-life was calculated as ln(2) divided by the decay rate after quality filtering; the exponential lifetime reported by Chen et al. was converted to half-life by multiplication by ln(2). (**C** and **G**) Absolute protein-synthesis rates from Li et al. (11), expressed as molecules per generation in MOPS complete or MOPS minimal medium; bracketed low-confidence measurements supported by fewer than 128 reads were excluded. (**D** and **H**) Protein abundance from Schmidt et al. (12), expressed as copies per cell during growth in LB or M9 glucose medium and after 1 or 3 days in stationary phase. Where multiple protein entries mapped to the same gene, copies per cell were summed. All x-axes are logarithmic, and the horizontal line at a cumulative fraction of 0.5 indicates the median. Each inset reports the fold change (FC), nominal two-sided Wilcoxon rank-sum P value and numbers of AR2/control genes retained for that condition. Fold changes are ratios of the AR2 and control geometric means; each dataset and condition was tested separately without multiple-testing adjustment.

#### Supplementary Figure 5

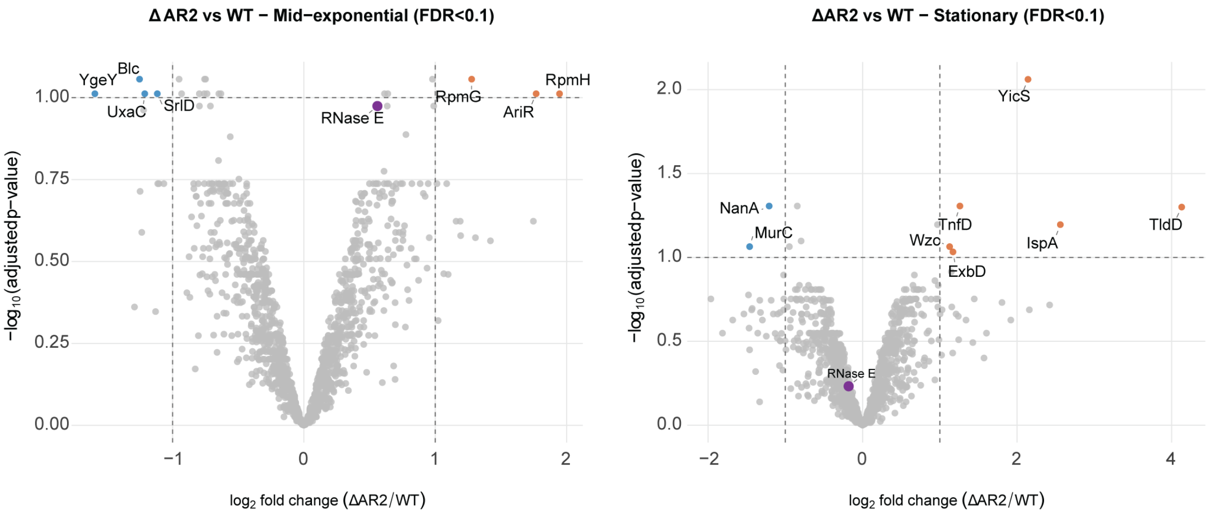


**SI Figure 5. Differential protein abundance following deletion of the RNase E AR2 domain.**

Volcano plots compare label-free protein abundance in ΔAR2 and wild-type cells during (**A**) mid-exponential and (**B**) stationary phase. MaxQuant LFQ intensities (13, 14) from four biological replicates per genotype and growth phase were $\log_2$-transformed, with zero intensities treated as missing values. Differential abundance was analysed using linear models with empirical Bayes moderation in limma (15). Positive log2 fold change indicates greater abundance in ΔAR2 than wild type. FDR was calculated using the Benjamini–Hochberg procedure. Dashed vertical lines indicate $\log_2$ fold changes of −1 and +1, and the dashed horizontal line indicates a false-discovery rate (FDR) of 0.10. Proteins satisfying both FDR < 0.10 and |log2 fold change| ≥ 1 are coloured and labelled: blue denotes lower abundance and orange denotes higher abundance in ΔAR2. Grey points do not satisfy both thresholds. RNase E (*rne*) is highlighted in purple (mid-exponential: $\log_2$ fold change = 0.56, FDR = 0.106; stationary: $\log_2$ fold change = −0.18, FDR = 0.586).
