## Supplementary Methods for "The intrinsically disordered AR2 domain of RNase E binds mRNA translation initiation regions"

### EXPERIMENTAL METHODS

#### Bacterial strains and general culture conditions

The bacterial strains, plasmids and oligonucleotides used in this study are listed in Supplementary Tables 1-3. E. coli str. K-12 substr. MG1655 or DH5α were routinely cultured at 37°C on solid or in liquid Luria-Bertani (LB) media, unless otherwise stated. Antibiotics were routinely used in this study to select for plasmids at 100 µg/mL ampicillin, 50 µg/mL kanamycin or 50 µg/mL spectinomycin, unless otherwise specified. All bacterial strains were stored at −80°C as stationary phase cultures with 16% (v/v) glycerol.

#### Bacterial strain construction

Modifications to the chromosomal copy of RNase E in E. coli MG1655 were performed using CRISPR-Cas9 and the pCas/pTarget dual-vector system [1]. These modifications included: insertion of a C-terminal $His_6$–TEV–3×FLAG (HTF) tag into full-length RNase E (constructing RNase E-HTF); separation of the AR2 microdomain from the RNase E-HTF strain by the insertion of a second TEV cleavage site upstream at the Val744 codon allowing dual-affinity purification of the AR2 CTD fragment by the downstream HTF tag (AR2-HTF); and deletion of the AR2 CTD-containing region from wild-type MG1655 (ΔAR2). Briefly, selection of pCas in electrocompetent MG1655 was performed at 30°C on solid LB supplemented with 50 µg/mL kanamycin. MG1655 pCas was then cultured to an OD600 0.10, induced with 50 mM L-arabinose for 1 h and made electrocompetent. The pTarget vector and linear double-stranded DNA products containing either the HTF, TEV or AR2 deletion, at least 250-nt of flanking homology regions and the appropriate guide (g)RNA were co-transformed into MG1655 pCas and selected at 30°C on solid LB supplemented with 50 µg/mL kanamycin and 50 µg/mL spectinomycin. The co-transformed linear DNA products were either generated by PCR (RNase E-HTF strain) or synthesised as double-stranded gBlocks (AR2-HTF and ΔAR2 strains) (IDT) and are described in Supplementary Table 3. Transformants were screened using

PCR with primers flanking the appropriate insertion or deletion region in the chromosome (Supplementary Table 3) and successful colonies were cured of pTarget by overnight incubation at 30°C in liquid LB supplemented with 0.5 mM IPTG. The pCas plasmid was cured by successive overnight passages at 40°C in liquid LB containing no antibiotics. Loss of the pCas and pTarget vectors were confirmed in MG1655 RNase E-HTF, AR2-HTF and ΔAR2 strains by kanamycin and spectinomycin sensitivity, and plasmid-specific PCR (Supplementary Table 3).

**Split-CRAC and CLASH**

CLASH was performed as previously described in E. coli [2], with updates from the protocol that was applied to S. aureus utilising the endoribonuclease RNase III [3, 4]. MG1655 WT (untagged), RNase E-HTF and AR2-HTF strains were used to inoculate 50 mL of pre-warmed liquid LB and grown at 37°C to an OD600 2.0 (early stationary phase) with 180 rpm shaking. Cultures were crosslinked with 1800 mJ dosage of UV-C irradiation (Vari-X-Link, UV03) and immediately harvested by centrifugation (4,500 g for 15 min). Each culture condition was performed in triplicate (n=3). To each bacterial cell pellet, 1 mL of lysis buffer (50 mM Tris-HCl [pH 7.8], 1.5 mM $MgCl_2$, 150 mM NaCl, 0.1% NP-40, 5 mM β-mercaptoethanol and 1 tablet of "cOmplete" EDTA-free protease inhibitor [Roche] per 50 mL of buffer) and 1 V of 0.1 mm zirconia beads were added and vortexed for 2 × 40 s intervals using the FastPrep-24 5G (MP Biomedicals). Dry ice was used to ensure cell pellets remained chilled during intervals. Cell debris was centrifuged (4,500 g for 10 min) and the clarified lysate was transferred to 1.5 mL microcentrifuge tubes and further clarified at 16,000 g for 20 min. Supernatants were added to 100 µL equilibrated M2 anti-FLAG resin (Sigma) and incubated at 4°C with gentle rotation for 16 h. The resin was then washed once in 5 mL TNM1000 buffer (50 mM Tris-HCl [pH 7.8], 1 M NaCl, 0.1% NP-40, 5 mM β-mercaptoethanol) and twice in 5 mL TNM150 buffer (50 mM Tris-HCl pH 7.8, 150 mM NaCl, 0.1% NP-40, 5 mM β-mercaptoethanol). The protein-bound resin was resuspended in 500 µL of TNM150 buffer and incubated with 50 U of GST.TEV protease at 24°C with rotation for 2 h. Eluates were collected by filtration through a Bio-spin chromatography column (Bio-Rad) and incubated with 0.15 U of RNace-IT (Agilent) for 5 min at 20°C. The digestion was then stopped by the addition of 0.4 g guanidine-HCl, 300 mM NaCl and 10 mM imidazole. Eluates were added to 200 µL magnetic Ni-NTA resin (Thermo) equilibrated with wash buffer 1 (4 M guanidine-HCl, 50 mM Tris-HCl [pH 7.8], 300 mM NaCl, 0.1% NP-40, 5 mM β-mercaptoethanol) and incubated at 4°C with gentle rotation for 16 h. Ni-NTA resin was washed once with 1 mL wash buffer 1 and four times with 1 mL PNK buffer

(50 mM Tris-HCl [pH 7.8], 10 mM MgCl2, 0.1% NP-40 and 5 mM β-mercaptoethanol). The resin was resuspended in 80 μL NP-PNK buffer (50 mM Tris-HCl [pH 7.8], 10 mM MgCl2 and 5 mM β-mercaptoethanol) containing 8 U of alkaline phosphatase (Promega) and 80 U of recombinant RNasIN (Promega) and incubated at 37°C with rotation for 1 h. The magnetic Ni-NTA resin was washed once with 1 mL wash buffer 1 and four times with 1 mL PNK buffer, and resuspended in 80 μL NP-PNK buffer containing 20 U of T4 polynucleotide kinase (NEB), 20 U of RNasIN (Promega) and 30 μCi [γ-32P]ATP and incubated at 37°C with rotation for 45 min. Cold 1 mM ATP was spiked in and the reaction allowed to proceed for another 15 min to ensure all 5′ end phosphates are attached. Ni-NTA resin was again washed with 1 mL wash buffer 1 and four times with 1 mL PNK buffer and then resuspended in 80 μL of NP-PNK buffer containing 40 U of T4 RNA ligase I (NEB), 1 mM ATP, 20 U of RNasIN (Promega) and 100 pM of an L5 index barcoded 5′ linker (IDT) (Supplementary Table 3). The ligation reaction was incubated at 16°C with gentle rotation for 16 h. The magnetic Ni-NTA resin was washed with wash buffer 1 followed by PNK buffer and resuspended in 60 μL NP-PNK buffer containing 40 U of T4 RNA ligase I (NEB), 20 U of RNasIN (Promega) and 80 pM of a 3′ linker (IDT) (Supplementary Table 3). The ligation reaction was incubated at 16°C with gentle rotation for 16 h. Ni-NTA resin was washed once with 1 mL wash buffer 1 and four times with 1 mL wash buffer 2 (50 mM Tris-HCl [pH 7.8], 50 mM NaCl, 0.1% NP-40, 5 mM β-mercaptoethanol and 10 mM imidazole). $^{32}$P-radiolabelled full-length RNase E-RNA and AR2-RNA complexes were eluted in 200 μL of elution buffer (50 mM Tris-HCl [pH 7.8], 50 mM NaCl, 0.1% NP-40, 5 mM β-mercaptoethanol and 250 mM imidazole) at 8°C with rotation for two 15-min repeats and resolved on a NuPAGE 4-12% gradient Bis-Tris PAGE (Invitrogen) run at 4°C in 1x NuPAGE running buffer (Invitrogen) for 1.5 h at 125 V. $^{32}$P-radiolabelled full-length RNase E-RNA and AR2-RNA complexes were visualised by autoradiography, gel excised and recovered by incubating the fragmented gel pieces in 600 μL of wash buffer 3 (50 mM Tris-HCl [pH 7.8], 50 mM NaCl, 0.1% NP-40, 5 mM β-mercaptoethanol, 1% SDS and 5 mM EDTA) and 100 μg of proteinase K at 55°C for 2 h. Fragments of the gel were similarly excised for the wild-type (untagged) corresponding to the expected sizes of full-length RNase E and AR2, and processed identically. Gel pieces were removed by filtration through a Bio-Spin chromatography column (Bio-Rad) and RNA eluates were phenol:chloroform extracted and ethanol precipitated overnight at -80°C. Reverse transcription was performed using the RT_PE_reverse oligonucleotide (Supplementary Table 3) and SuperScript IV (Invitrogen). cDNA was incubated with RNase H (Invitrogen) at 37°C for 20 min and amplified using Phusion polymerase (NEB). A combination of either 10 pM BC1, BC2 or BC3 oligonucleotides

were used with 10 pM of the P5 primer to alternate the barcoded indexes (Supplementary Table 3). The cDNA libraries were amplified for 24 cycles with 2 µL of cDNA used as template. Libraries were separated on a 1.5% metaphor agarose gel and amplicons excised and purified using the MiniElute gel extraction kit (Qiagen). Libraries were sequenced on an Illumina NovaSeq6000 platform generating 150 bp paired-end reads (PE150) (Novogene, Singapore).

**Analysis of RNA–protein and RNA–RNA datasets**

UV-crosslinked reads were processed using the snakemake pipeline, Hyb-CRAC-R as described previously [3] which incorporates pyCRAC [5] for mapping CRAC reads and hyb [6] for mapping CLASH reads. The pipeline also includes scripts for calculating the statistical significance of RNA-RNA hybrids. Reads were mapped to the E. coli str. K-12 substr. MG1655 genome (NCBI NC_000913.3). Additional sequence features including 5′ and 3′ UTR annotations were obtained from RegulonDB [7]. pyCRAC [5] was used to count reads at specific sequence features. Reads were normalised to the total number of mapped reads in the library. Custom scripts were used to count reads and deletions at the start and end of genomic features. Hybrid counts in Figure 1E represent the total number of interactions across biological replicates (not filtered for statistical significance).

**RNase E motif analysis**

High confidence sites of direct contact with the AR2 domain were identified using contact-dependent deletions. Sites with ≥10 deletions were extracted with 50-nt of flanking sequence. Overlapping sites were merged using BEDTools merge [8] to remove duplicate sequences. Motifs were extracted using MEME-ChIP (MEME Suite) [9] with maximum width set at 6 or 10-nt. To examine AR2 binding to motifs, FIMO (MEME Suite) [10] was used to identify matching sites in the genome (FDR < 0.0001, --thresh). AR2 binding was mapped at motifs using custom awk scripts. For nucleotide counts at deletion sites, the unmerged deletion sites were used to preserve the spacing between the 5′ nucleotide and crosslinking site (merged reads use the longest 5′ and 3′ position). Nucleotide counts were determined using custom python scripts.

**Protein extracts and Western blot analysis**

Full-length RNase E and the AR2 domain from the MG1655 RNase E-HTF and AR2-HTF strains were dual-affinity purified as described for split-CRAC analyses. Purified proteins were run on a NuPAGE 4-12% gradient Bis-Tris PAGE gel (Invitrogen) in 1x NuPAGE MOPS running buffer (Invitrogen) for 1 h at 200 V. Gel excisions corresponding to bands of the RNase

E, the AR2 domain and a negative control (empty lane) were used for LC-MS/MS analyses to confirm the correct amino acids. Proteins separated from PAGE analyses were transferred onto a 0.2 µm nitrocellulose membrane (Thermo) in NuPAGE transfer buffer (Invitrogen) for 1.5 h at 30 V. The membrane was rinsed in water and washed 3x for 5 min in TBST (1x Tris-buffered saline and 0.1% Tween-20). Membranes were blocked for 1 h at room temperature in 5% skim milk and washed 3 x for 5 min in TBST. Blocked membranes were incubated in a 1/2000 dilution of 6x-His tag monoclonal antibody (Invitrogen) in 5% skim milk at 4°C with gentle rotation for 16 h. Membranes were washed 3x for 5 min in TBST and antibody detected using the Pierce ECL substrate blotting system (Invitrogen). Membranes were visualised using a LAS-3000 imaging system (Fujifilm, Japan) with automatic exposure settings.

**Expression and purification of RNase E(689-850) and RNase E-RhlB-enolase complex**

The RNase E(689–850)–RhlB–enolase complex was purified using a protocol adapted from Bruce *et al.* [11]. *E. coli* T7 Express BL21(DE3) cells were co-transformed with pRSFDuet-1 encoding C-terminally $His_6$-tagged RNase E(689–850) and untagged RhlB, and pET11a encoding untagged enolase. Cultures were grown in LB supplemented with 50 µg $mL^{-1}$ kanamycin and 100 µg $mL^{-1}$ ampicillin at 37°C with shaking at 200 rpm. At $OD_{600} \approx 0.5$, protein expression was induced with 1 mM IPTG and cultures were incubated overnight at 18°C. Cells from 4 L of culture were harvested by centrifugation and resuspended in 100 mL lysis buffer containing 50 mM Tris-HCl (pH 7.5), 250 mM KCl, 500 mM NaCl, 10 mM $MgCl_2$, 10% (v/v) glycerol, 1 mM DTT and protease inhibitor. Cells were lysed by sonication on ice (2 s on/2 s off, 50% amplitude, 8 min) and lysates were clarified by centrifugation at 15,000 × *g* for 30 min at 4°C.

The clarified lysate was incubated for 1 h at 4°C with Ni Sepharose 6 Fast Flow resin pre-equilibrated in Buffer A (50 mM Tris-HCl [pH 7.5], 500 mM NaCl, 250 mM KCl, 10 mM $MgCl_2$ and 1 mM DTT). To remove contaminating nucleic acids, the resins were washed with approximately 30 mL Buffer A containing 100 mM urea and subsequently re-equilibrated with Buffer A. The RNase E(689–850)–RhlB–enolase complex was eluted with Buffer A containing 250 mM imidazole. Elution fractions were analysed by 12% SDS-PAGE and fractions containing RNase E(689–850), RhlB and enolase were pooled and dialysed at 4°C against dialysis buffer (50 mM Tris-HCl (pH 7.5), 200 mM NaCl, 200 mM KCl, 10 mM $MgCl_2$ and 1 mM DTT).

RNase E(689–850) was subsequently isolated from the ternary complex by rebinding the complex to fresh Ni Sepharose resin through the C-terminal $His_6$ tag. The resins were washed with Buffer A followed by a denaturing wash containing 50 mM Tris-HCl (pH 7.5), 200 mM KCl, 400 mM NaCl, 8 M urea, 10 mM $MgCl_2$ and 1 mM DTT to dissociate untagged RhlB and enolase, as previously described [11]. Following re-equilibration with Buffer A, $His_6$-tagged RNase E(689–850) was eluted with Buffer A containing 250 mM imidazole. Fractions containing purified RNase E(689–850) were identified by 12% SDS-PAGE, pooled and dialysed as described above. Purified proteins were aliquoted, flash-frozen in liquid nitrogen and stored at −80°C.

**Electrophoretic mobility-shift assays**

Electrophoretic mobility-shift assays were adapted from Bruce et al. [11]. The 66-nt wild-type lpp translation-initiation-region RNA and the corresponding motif-substitution mutant (lpp-5G) were generated by T7 in vitro transcription from PCR-amplified templates. RNAs were dephosphorylated using Quick CIP (New England Biolabs), purified using an RNA Clean & Concentrator kit (Zymo Research), and 5′-end-labelled with [$\gamma$-$^{32}$P]ATP using T4 polynucleotide kinase (New England Biolabs).

Binding reactions contained 100 fmol labelled RNA and the indicated concentration of isolated AR2 (0–2,000 nM) or AR2–RhlB–enolase complex (0–200 nM) in 10 µL GF buffer (20 mM Tris–HCl, pH 8.0, 150 mM NaCl and 5 mM $MgCl_2$). Reactions were incubated at 30°C for 30 min and supplemented with 2 µL loading dye containing 20% glycerol, bromophenol blue and xylene cyanol. Complexes were resolved by 5% native polyacrylamide-gel electrophoresis and detected by phosphorimaging. Bound and unbound RNA signals were quantified by densitometry, and the bound fraction was calculated as bound RNA divided by the sum of bound and unbound RNA. Binding curves were fitted in GraphPad Prism using nonlinear regression with the “specific binding with Hill slope” model, from which apparent dissociation constants ($K_d$) were estimated.

**Label-free quantitation of the proteome using LC-MS/MS**

*E. coli* MG1655 wild-type and ΔAR2 strains were cultured in LB at 37°C with shaking at 220 rpm and harvested during mid-exponential and stationary growth phases. Four independent biological replicates were prepared for each strain and growth phase. Cell pellets were resuspended in lysis buffer containing 1% (w/v) sodium deoxycholate (SDC) and 100 mM Tris-HCl (pH 8.0) and mechanically lysed with 0.1-mm zirconia beads. Lysates were clarified by centrifugation and protein concentrations were normalised. For each sample, 50 μg of total protein was submitted to the UNSW Bioanalytical Mass Spectrometry Facility (BMSF) for tryptic digestion and LC–MS/MS analysis using an Orbitrap Exploris 480 mass spectrometer.

Raw LC-MS/MS data were processed using MaxQuant v2.7.5.0 [12, 13] against the UniProt *E. coli* K-12 reference proteome (UP000000625). Searches used Trypsin/P with up to two missed cleavages, with carbamidomethylation (C) specified as a fixed modification and oxidation (M) and protein N-terminal acetylation as variable modifications. Peptide-spectrum matches and protein identifications were controlled at 1% FDR, and label-free quantification was performed with match-between-runs enabled. LFQ intensities from four biological replicates per genotype and growth phase were $\log_2$-transformed, with zero intensities treated as missing values.

**MaxQuant LFQ preprocessing and differential protein abundance**

The MaxQuant proteinGroups output was used for LFQ-based analysis. Entries lacking a gene name were removed, LFQ intensities of zero were treated as missing values, and intensities were $\log_2$-transformed. Duplicate protein-group rows sharing a gene name were averaged within each sample. Mid-exponential and stationary samples were analysed separately. For SI Figure 5, a protein was retained within a phase when it was observed in at least two of four replicates in both wild-type and ΔAR2 cells; missing values were not imputed. Differential abundance was estimated with limma linear models and empirical Bayes moderation [14]. Positive $\log_2$ fold change denotes higher abundance in ΔAR2 than wild type. P values were adjusted across proteins within each phase using the Benjamini–Hochberg procedure. Volcano-plot classifications required FDR < 0.10 and $|\log_2 \text{ fold change}| \geq 1$. RNase E was highlighted independently of these thresholds.

**RNase E abundance analysis**

For the RNase E abundance plot in main Figure 5B, the $\log_2$ LFQ matrix was median-normalised separately within each sample. The rne values were extracted and centred by subtracting the mean abundance of the mid-exponential wild-type group, so that the reference-group mean equalled zero. Individual biological replicates are shown together with the group mean and standard deviation. Wild-type and ΔAR2 values were compared independently within each growth phase using two-sided Mann–Whitney U tests. Centring on the mid-exponential wild-type mean changes the plotted origin but not the within-phase group comparison.

### DEFINITION OF AR2 AND CONTROL GENE SETS

The stringent AR2-crosslinked gene set was constructed from the union of AR2 peaks detected in one or more biological replicate. Genes were retained when an AR2 crosslinking peak occurred within 20 nt of the annotated translation start site, the region contained more than 10 contact-dependent deletions per nucleotide, and the associated sequence contained a statistically significant match to the extended A-rich AR2 motif (FDR < 0.001). This produced 107 unique genes.

For analyses of sequence and translation-initiation features, the comparison set comprised genes annotated by RegulonDB as the first gene in an operon. This control was selected so that the comparison genes were expected to possess their own translation-initiation regions. The initial list contained 1,595 unique genes. Genes belonging to the AR2 set were removed from the control before each analysis, leaving 1,541 mutually exclusive control genes for analyses using the stringent AR2 set.

An additional motif-only set was used in SI Figure 4 to distinguish properties associated with AR2 crosslinking from those associated with occurrence of the motif alone. This set comprised 250 unique genes with a FIMO match to the extended A-rich motif within 20 nt of the annotated translation start site (FDR < 0.0001), without requiring an AR2 crosslink-dependent deletion. Eighty-eight motif-set genes were present in the first-in-operon list and were removed from that control, leaving 1,507 control genes. Sample sizes differ among panels because only genes with measurements in the relevant published dataset were included. Missing measurements were not imputed.

### FUNCTIONAL ENRICHMENT, SRP INTERACTION AND PROTEIN LOCALISATION (SI FIGURE 2)

#### Gene Ontology enrichment

Gene Ontology enrichment was performed using GOEnrichment on the Galaxy platform (usegalaxy.org) [15, 16]. The stringent AR2 gene list was analysed separately against the Biological Process, Molecular Function and Cellular Component ontologies using the Escherichia coli annotation population implemented by the tool. Enrichment results and multiple-testing-adjusted q values were taken from the GOEnrichment output. No ontology term met an FDR threshold of 0.05. The highest-ranking Biological Process term was "negative regulation of translational elongation" (GO:0045900; raw $P = 2.2 \times 10^{-4}$, $q = 0.0679$; genes raiA and ettA); no terms were returned for the Molecular Function or Cellular Component ontologies.

#### Signal-recognition-particle interaction

SRP interaction measurements were obtained from the E. coli SRP interactome reported by Schibich et al. [17]. The continuous SRP-interaction measure was calculated as SRP immunoprecipitation reads per million divided by total-RNA reads per million. Duplicate gene-symbol entries in the source workbook were averaged to produce one measurement per gene. Historical and current gene symbols were reconciled using exact symbols and unambiguous aliases. Genes absent from the SRPome were treated as unmeasured and were not assigned a value of zero.

The primary background comprised every other gene with a valid ratio in the same SRPome experiment, excluding AR2 genes. This yielded 84 AR2-crosslinked genes and 2,281 background genes. Ratios were displayed on a log2-spaced axis, and the distributions were compared using a two-sided Wilcoxon rank-sum test. Cliff's delta was calculated as a rank-based effect-size measure, with positive values indicating greater SRP interaction among AR2 genes.

A binary SRP-client membership was taken from the published SRP-client classification, corresponding to the reported SRP-enrichment criterion of greater than 2.0. The proportions of classified clients in the AR2 and background sets were compared using a two-sided Fisher's exact test, and an odds ratio with 95% confidence interval was reported. The first-in-operon genes represented in the SRPome, after exclusion of AR2 genes, were analysed separately as a sensitivity comparison.

#### Predicted protein localisation

Subcellular-localisation predictions for E. coli K-12 MG1655 (RefSeq NC_000913) were downloaded from cPSORTdb/PSORTdb [18, 19]. The downloaded database release is PSORTdb 4.0, whereas the computational predictions reported in the Final_Localization field were generated by the PSORTb 3.0 algorithm. RefSeq protein accessions were mapped to gene symbols using the corresponding E. coli genome annotation, and historical gene symbols were reconciled using unambiguous aliases.

Of 105 AR2 genes mapped to the PSORTb export, 84 had a resolved final localisation and 21 were classified as unknown. Resolved predictions were assigned to cytoplasmic, cytoplasmic-membrane, periplasmic, outer-membrane or extracellular categories. The primary comparison used all other genes with resolved predictions in the same PSORTb export (n = 3,181). The overall distribution of localisation categories was tested using a two-sided Fisher's exact test on the 2 × 5 contingency table. Each localisation was then tested separately against all other resolved localisations using a two-sided Fisher's exact test. Odds ratios and 95% confidence intervals were calculated, and P values were adjusted across the five category-specific tests using the Benjamini–Hochberg procedure. A separate Fisher's exact test compared the frequency of unknown predictions between AR2 and all other mapped genes. A first-in-operon background, excluding AR2 genes, was also analysed as a sensitivity comparison.

### TRANSLATION-INITIATION FEATURES (SI FIGURE 3)

#### Annotated 5′-UTR length

Strand-aware 5′-UTR annotations and coordinates were obtained from RegulonDB v12.0 [7]. All distinct annotated 5′ UTRs were retained, including alternative leaders associated with different transcription start sites. Consequently, the unit of analysis and the sample size in this panel are annotated UTR records rather than unique genes, and a gene can contribute more than one record. UTR lengths were displayed on a logarithmic axis and compared between the AR2 and first-in-operon sets using a two-sided Wilcoxon rank-sum test. Rank-biserial correlation was reported as an effect-size measure.

#### Predicted translation-initiation rate

Translation-initiation rates were predicted with RBS Calculator v1.0 [20], using the sequence associated with each annotated start site and the annotated start-codon position. The legacy workflow used Python 2.7.18, ViennaRNA and NUPACK 3.0.6. Where alternative UTR annotations shared a gene name, the gene-named calculator output yielded one retained result per gene. Successful predictions were obtained for 62 AR2 genes and 690 first-in-operon control genes. Predicted rates were plotted as log10 values and compared using a two-sided Wilcoxon rank-sum test.

#### Nucleotide composition surrounding the translation start

Strand-oriented sequences spanning positions −60 to +29 relative to the first nucleotide of the annotated start codon were extracted for each gene. The 26-nt region from −20 to +5, inclusive, was retained for analysis. DNA thymidine characters were converted to uridine, and the percentages of A, U, C and G were calculated separately for each gene. All 107 AR2 genes were retained, including five genes with annotated GUG start codons. After removal of 54 overlapping AR2 genes, the first-in-operon comparison contained 1,541 genes.

For each nucleotide, the AR2 and control distributions were compared using a two-sided Wilcoxon rank-sum test without continuity correction. P values were adjusted across the four nucleotide comparisons using the Benjamini–Hochberg procedure. Rank-biserial correlation was calculated from the Mann–Whitney statistic as $2U/(n\_AR2 \times n\_control) - 1$, with positive values indicating higher nucleotide content in AR2-crosslinked genes. Boxplots show medians and interquartile ranges, with whiskers extending to the most extreme value within 1.5 times the interquartile range. Outlying observations were included in all calculations and tests but were not plotted individually.

**Shine–Dalgarno and start-codon sequence logos**

For the broad Shine–Dalgarno comparison, all 7-nt windows within the 25 nt upstream of each start codon were scored for hybridisation to the 16S rRNA anti-Shine–Dalgarno sequence ACCUCCU using UNAFold hybrid-min. The window with the most favourable predicted hybridisation energy was selected, and a 25-nt sequence beginning at the 5′ boundary of the predicted Shine–Dalgarno match was extracted. This produced 107 AR2 sequences and 2,383 non-AR2 sequences for SI Figure 3D and E.

For SI Figure 3F–I, the mRNA nucleotides predicted by RBS Calculator v1.0 to pair with 16S rRNA were extracted from the predicted mRNA–rRNA structure. Sequences were aligned either to the 5′ boundary of this predicted paired span or to the annotated start codon. Alternative records were deduplicated by gene, producing 62 AR2 and 690 first-in-operon control sequences. Sequence logos were generated with WebLogo 3.9.0 [21] using RNA lettering and the standard nucleotide colour scheme. Total stack height represents positional information content in bits, and relative letter height represents nucleotide frequency at each position.

### mRNA ABUNDANCE, mRNA HALF-LIFE, PROTEIN SYNTHESIS AND PROTEIN ABUNDANCE (SI FIGURE 4)

The stringent AR2-crosslinked set and the motif-only set were analysed separately using the same processing and statistical workflow. In each analysis, overlapping genes were removed from the first-in-operon control before the published datasets were joined by exact gene symbol. Only finite, positive measurements were retained. Missing values were not imputed, and the number of genes therefore varies by dataset and condition.

**mRNA abundance and half-life**

mRNA abundance and decay measurements were obtained from untreated wild-type cells in Moffitt et al. [22]. For mRNA abundance, the reported abundance at 0 min was averaged across the two untreated wild-type replicates to produce one positive value per gene.

For half-life estimation, decay rates from untreated wild-type samples were converted to half-lives as ln(2)/k, where k is the reported decay rate in $min^{-1}$. The reported 95% confidence interval was parsed for each fitted decay rate, and one quarter of its width was used as an approximation of the fit error. Measurements were retained when the decay rate was finite and positive, the fit error was less than half of the estimated decay rate, and the calculated half-life was no greater than 20 min. Passing replicate measurements were averaged by gene.

An independent half-life dataset was obtained from Chen et al. [23]. The reported average lifetime is the exponential time constant and was converted to half-life by multiplication by ln(2). Only exact, unambiguous gene-symbol matches were retained. Results from Moffitt et al. [22] and Chen et al. [23] were analysed separately and were not pooled.

**Absolute protein synthesis**

Absolute protein-synthesis measurements were obtained from Table S1 of Li et al. [24]. Measurements from MOPS complete and MOPS minimal media were analysed separately and are expressed as protein molecules produced per generation. Values enclosed in square brackets in the source table, indicating estimates supported by fewer than 128 sequencing reads, were excluded. These measurements represent absolute protein synthesis and were not normalised to mRNA abundance; they therefore do not measure translation efficiency per transcript.

**Protein abundance**

Protein copy numbers were obtained from the final combined estimates in Table S6 of Schmidt et al. [25]. Measurements from LB, M9 glucose, stationary phase day 1 and stationary phase day 3 were analysed separately. Where multiple protein or UniProt entries mapped to the same gene, copies per cell were summed to produce one gene-level value for each condition.

**Statistical analysis and visualisation**

Empirical cumulative distribution functions were plotted using the untransformed positive measurements with logarithmically scaled x axes. For each dataset and condition, the AR2 and control distributions were compared using a two-sided Wilcoxon rank-sum test with the asymptotic approximation. Fold changes were calculated as the ratio of the geometric mean in the AR2 set to the geometric mean in the corresponding control. The P values displayed in SI Figure 4 are nominal and were not adjusted for the number of datasets or conditions examined. Each published dataset and growth condition was treated as a separate descriptive comparison.

All analysis scripts, source-data tables, gene-mapping audits, statistical summaries and R session information used to generate SI Figures 2–4 are retained with the corresponding figure-specific reproducibility files.

### DIFFERENTIAL PROTEIN ABUNDANCE (SI FIGURE 5)

The SI Figure 5 volcano plots use the phase-specific, no-imputation limma workflow described above. The mid-exponential analysis included 1,495 proteins; four were lower and three higher in ΔAR2 at FDR < 0.10 and $|\log_2$ fold change$| \geq 1$. The stationary-phase analysis included 1,260 proteins; two were lower and six higher in ΔAR2 at the same thresholds. RNase E did not meet the FDR threshold in either phase (mid-exponential: $\log_2$ fold change = 0.559, FDR = 0.106; stationary: $\log_2$ fold change = −0.180, FDR = 0.586).

### STATISTICAL ANALYSIS

The statistical test, directionality, multiple-testing procedure and significance threshold used for each experiment are stated in the relevant figure legend and, for SI Figures 2–5, in the analysis-specific sections above. Unless otherwise stated, tests were two-sided. No observations were excluded solely on the basis of being statistical outliers.

### DATA AVAILABILITY

Split-CRAC sequencing data have been deposited in NCBI GEO under accession GSE317719. The reviewer token for GSE317719 is cnonqmeihxujdcx. Mass spectrometry proteomics data generated in this study are available at the ProteomeXchange Consortium via the PRIDE partner repository with the dataset identifier PXD082328 (reviewer token: bU6SLAwOClFt).
